# Modeling of Severe Acute Respiratory Syndrome Coronavirus 2 (SARS-CoV-2) Proteins by Machine Learning and Physics-Based Refinement

**DOI:** 10.1101/2020.03.25.008904

**Authors:** Lim Heo, Michael Feig

## Abstract

Protein structures are crucial for understanding their biological activities. Since the outbreak of severe acute respiratory syndrome coronavirus 2 (SARS-CoV-2), there is an urgent need to understand the biological behavior of the virus and provide a basis for developing effective therapies. Since the proteome of the virus was determined, some of the protein structures could be determined experimentally, and others were predicted via template-based modeling approaches. However, tertiary structures for several proteins are still not available from experiment nor they could be accurately predicted by template-based modeling because of lack of close homolog structures. Previous efforts to predict structures for these proteins include efforts by DeepMind and the Zhang group via machine learning-based structure prediction methods, i.e. AlphaFold and C-I-TASSER. However, the predicted models vary greatly and have not yet been subjected to refinement. Here, we are reporting new predictions from our in-house structure prediction pipeline. The pipeline takes advantage of inter-residue contact predictions from trRosetta, a machine learning-based method. The predicted models were further improved by applying molecular dynamics simulation-based refinement. We also took the AlphaFold models and refined them by applying the same refinement method. Models based on our structure prediction pipeline and the refined AlphaFold models were analyzed and compared with the C-I-TASSER models. All of our models are available at https://github.com/feiglab/sars-cov-2-proteins.

## INTRODUCTION

Protein tertiary structures are essential for understanding their biological mechanisms. Such insight at the molecular level, allows those proteins to be exploited as therapeutic targets by identifying either already approved drug molecules that could be repurposed or discovering new drug candidates via computational methods such as virtual screening. Since the SARS-CoV-2 infection reached pandemic level since early 2020, there is now an urgent need for high-resolution structures of this virus. As protein sequences for the virus proteome were determined quickly^1^, some of the protein structures could be obtained experimentally. However, experimental structures of many proteins are still not available to date, leaving prediction via computational methods as the only alternative. SWISS-MODEL^2^ could predict tertiary structure models for a subset of proteins by relying on template-based modeling techniques. since many of the genes in the SARS-CoV-2 genome are close homologs to proteins in other organisms with known structures. However, for some of the proteins, template-based modeling is not possible because of lack of experimentally determined close homologs. Recently, the prediction of tertiary structures for proteins where no template structures are available, has been advanced significantly via novel machine learning methods^3^. This approach predicts inter-residue distances from multiple sequence alignment via deep learning. Using this approach, DeepMind applying the AlphaFold method^3^ to make predictions for six proteins, where close homolog structures are not available^4^. In addition, the Zhang group predicted models for the entire proteome^5^, including targets for which no homologs can be identified, by using the novel C-I-TASSER platform^6^, which is a combined method of contact-based and template-based modeling.

We also made predictions for those proteins which do not have close homolog structures and focused in particular on applying a high-resolution physics-based refinement protocol to improve the accuracy of machine-learning based models^7^. We followed two protocols: In the first, we generated initial machine learning-based models by using trRosetta^8^. In the second protocol we started from DeepMind’s AlphaFold models. Both sets of machine-learning based models were subjected to our latest molecular-dynamics based refinement protocol^9,10^ to maximize model accuracy. Here we compare the resulting models with each other and with the predictions from C-I-TASSER. A particular focus is on establishing, which structural aspects are conserved based on consensus from different approaches and where significant uncertainty remains in the accuracy of the computer-generated models. All of our predicted protein tertiary models are publicly available at https://github.com/feiglab/sars-cov-2-proteins.

## RESULTS

We predicted structures of 10 proteins from the SARS-CoV-2 proteome, as summarized in **Table 1**. Protein models were generated initially from inter-residue distance predictions rather than template-based modeling because of a lack of experimentally determined close homolog structures. The resulting models were then further refined by a molecular dynamics simulation-based refinement protocol to improve the physical realism at the atomistic level of the structures. We also applied the same refinement protocol to the models predicted by DeepMind’s AlphaFold method. The detailed procedures are described in **METHOD** section. We compared our models with the other available models, i.e. the original AlphaFold models and the predictions from the Zhang group. As shown in **Table 1**, our models and the models from the Zhang group provide more complete sequence coverage than the AlphaFold models.

**Table 1.** Summary of the modeled proteins and comparisons of predicted residues with other available models

| Protein name | RefSeq | FeigLab | AlphaFold | Zhang |
| --- | --- | --- | --- | --- |
| nsp2 | YP_009725298.1 | 1–638 | 1–345, 438–638 | 1–638 |
| nsp4 | YP_009725300.1 | 1–500 | 1–489 | 1–500 |
| nsp6 | YP_009725302.1 | 1–290 | 1–278 | 1–290 |
| PL-PRO (nsp3) | YP_009725299.1 | 1260–1945 | 1571–1927 | 1–1945 |
| ORF3a | YP_009724391.1 | 1–275 | 38–233 | 1–275 |
| Membrane glycoprotein | YP_009724393.1 | 1–222 | 11–203 | 1–222 |
| ORF6 | YP_009724394.1 | 1–61 | N/A | 1–61 |
| ORF8 | YP_009724396.1 | 1–121 | N/A | 1–121 |
| ORF10 | YP_009725255.1 | 1–38 | N/A | 1–38 |
| ORF7b | YP_009725296.1 | 1–43 | N/A | N/A |

When we applied our latest refinement protocol to AlphaFold protein models, the structure changes upon refinement were moderate, usually less than 2 Å in Cɑ-RMSD. (**Table 2**) The most significant changes occurred mostly in loops and the relative orientation between secondary structure elements. For example, in **Figure 1D**, a loop structure was changed, and the relative orientations of an α-helix and two β-strands were adjusted with respect to the other secondary structure elements. As another example, in **Figure 2D**, an α-helix was moved to improve hydrophobic packing and salt bridges between charged residues. We found earlier that machine learning-based models can be improved at the atomistic level by physics-based refinement and we expect that to the extent that the general features of the initial models are correct, refinement resulted in improved models.

**Table 2.** Structure change of AlphaFold models upon refinement measured in Cɑ-RMSD

| Protein name | Residues | Structure change upon refinement [Å] |
| --- | --- | --- |
| nsp2 | 1–345 | 1.50 |
|  | 438–638 | 1.86 |
| nsp4 | 1–273 | 1.79 |
|  | 274–399 | 1.90 |
|  | 400–489 | 0.80 |
| nsp6 | 1–278 | 2.01 |
| PL-PRO (nsp3) | 1571–1762 | 1.12 |
|  | 1763–1927 | 1.13 |
| ORF3a | 38–233 | 2.26 |
| Membrane glycoprotein | 11–203 | 1.27 |

**Figure 1.**
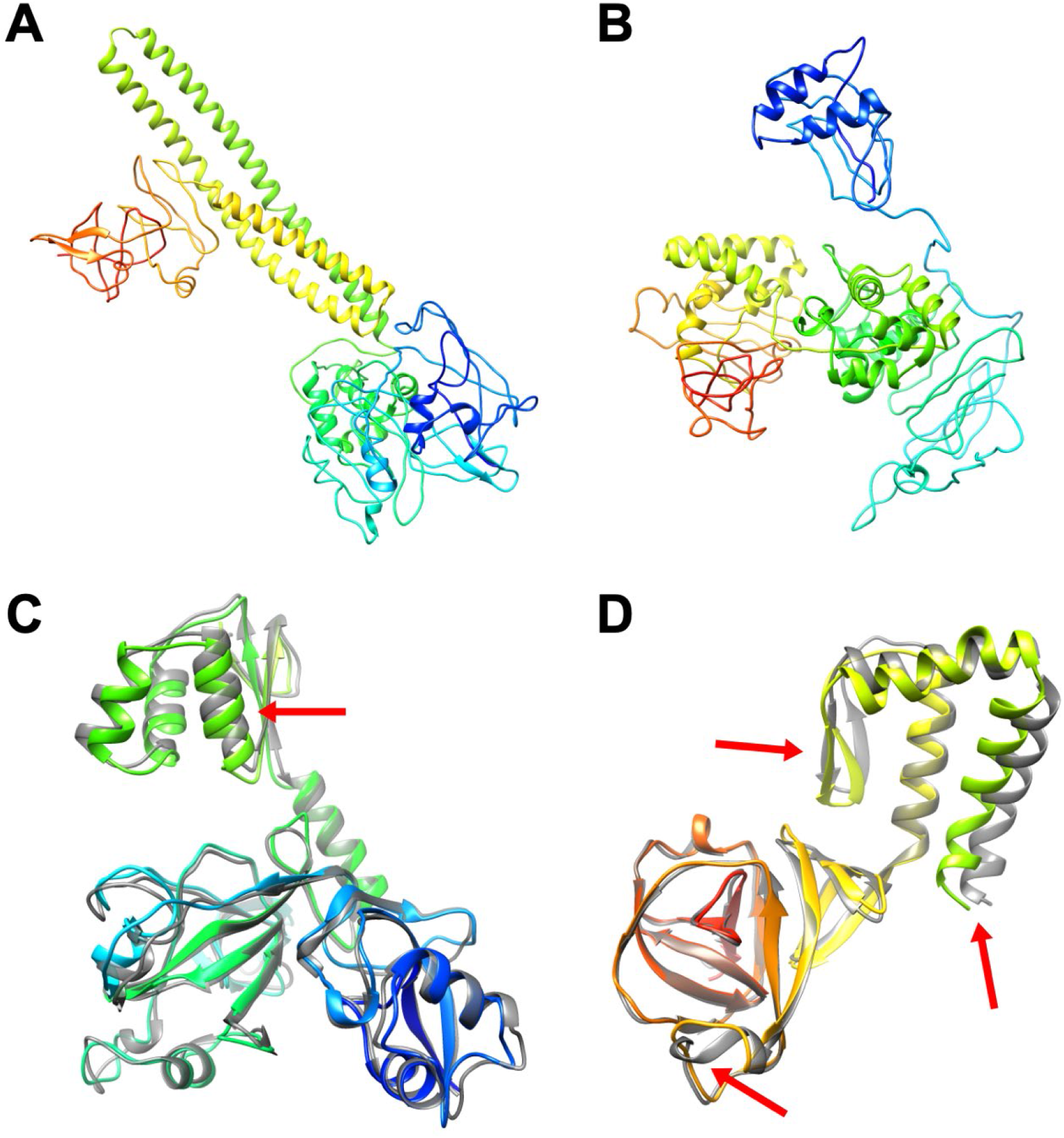
Protein models for nsp2: FeigLab (A), Zhang group (B), and AlphaFold models and their refined models for residues 1–345 (C) and 438–638 (D). Structures are shown in cartoon representation and colored in rainbow from blue (N-terminal) to red (C-terminal). (C and D) Refined AlphaFold models are shown in rainbow, while AlphaFold models are shown in grey. Significantly changed regions after refinement are indicated by red arrows.

**Figure 2.**
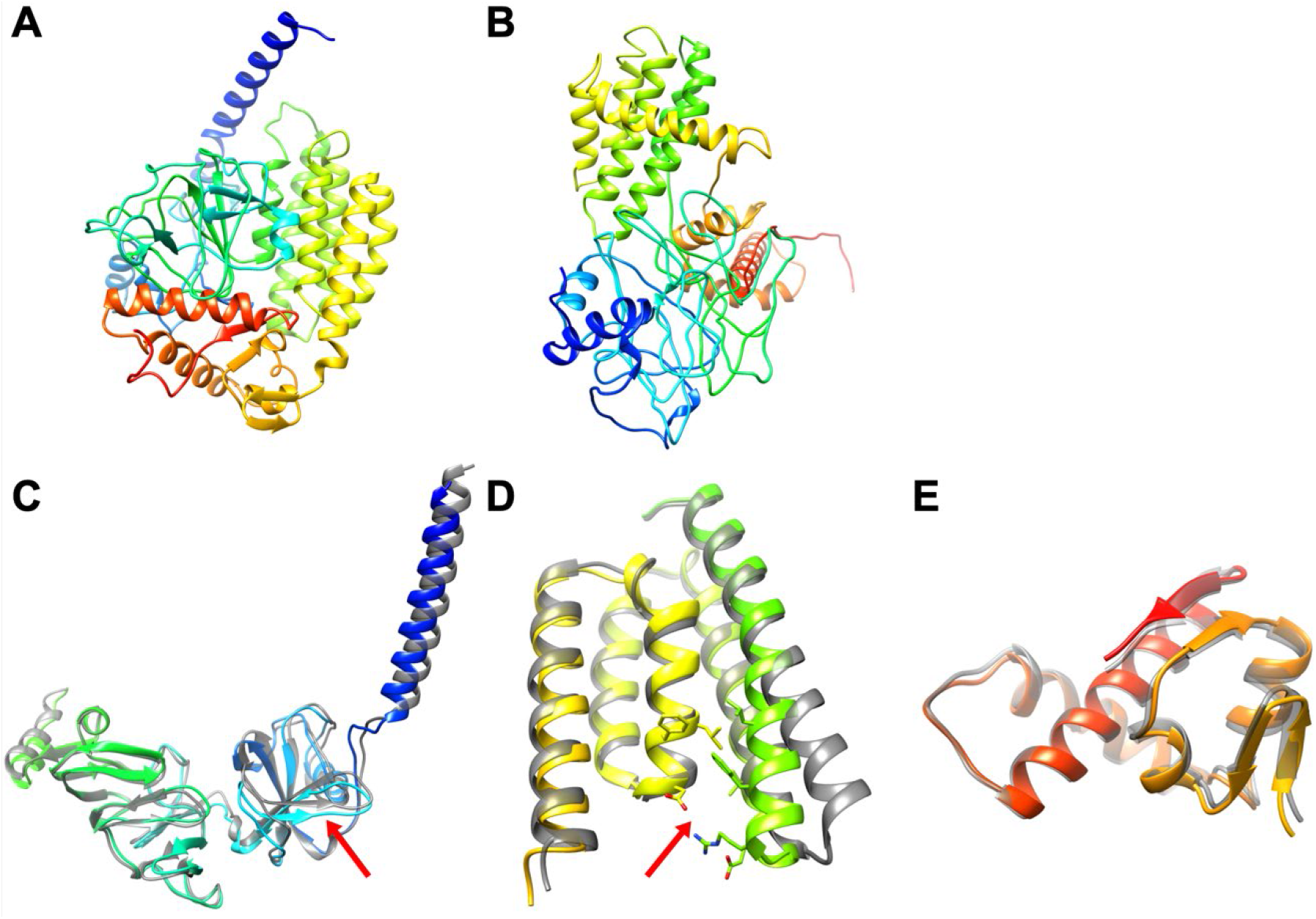
Protein models for nsp4: FeigLab (A), Zhang group (B), AlphaFold models and their refined models for residues 1–273 (C), 274–399 (D), and 400–489 (E). *See Figure 1*.

When comparing the models from our protocol with the (refined) AlphaFold models and the models from the Zhang group, we find that the resulting models do not reach a high degree of consensus for most of the modeled proteins (**Figures 1–7**). This may be expected as the modeled proteins were very difficult to predict. However, there is consensus on some of the proteins. For the domain of Papain-like proteinase (PL-PRO), residues 1763–1927, ours and AlphaFold’s model resemble each other with a Cɑ-RMSD of 2.96 Å. Moreover, for one of the domains of nsp4, residues 274–399, ours and AlphaFold’s model have the same topology, an α-helix bundle with the only significant difference being the orientation of the N-terminal helix (residues 274–309) that led to an overall difference between the structures of 8.27 Å in Cɑ-RMSD. Moreover, the membrane glycoprotein structure was also predicted with a similar topology between our protocol and AlphaFold with presumed trans-membrane N-terminal helices in a similar orientation and a globular C-terminal domain for residues 117–203 with a similar β-strand topology. The difference between our model and the AlphaFold prediction was 8.69 Å in Cɑ-RMSD mainly because of the orientation of two beta-strands (residues 117–133). Except for these cases, none of the structures shared a same topology.

**Figure 3.**
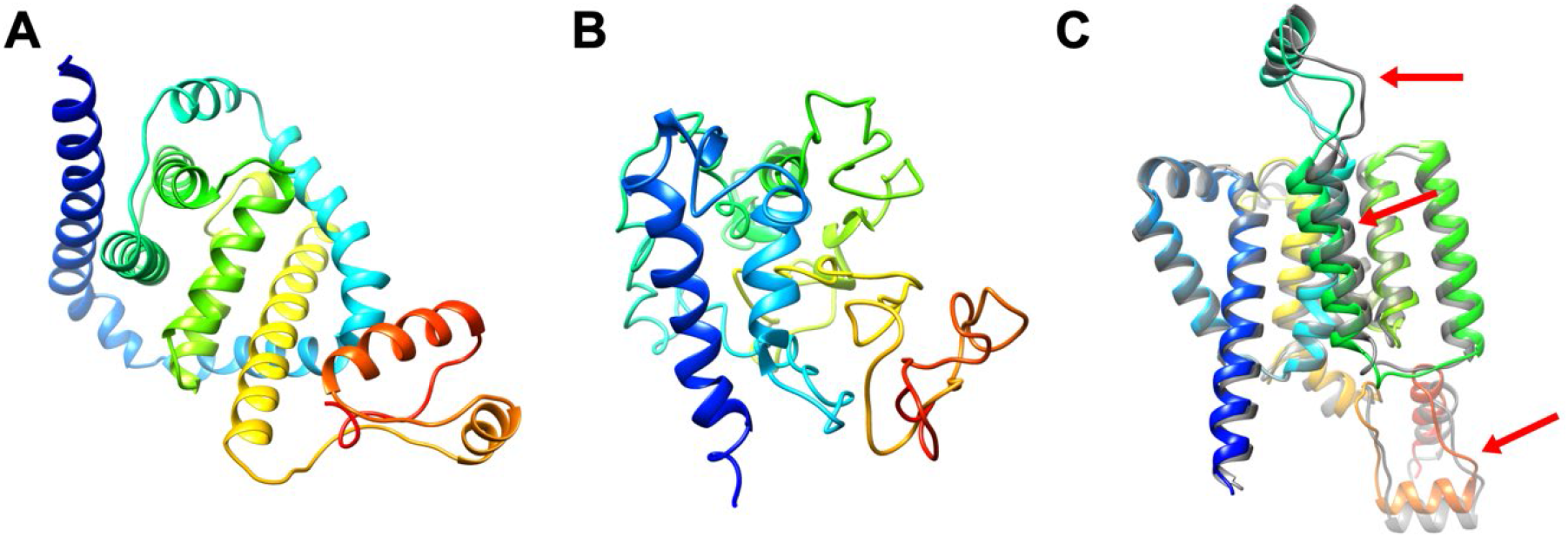
Protein models for nsp6: FeigLab (A), Zhang group (B), AlphaFold model and its refined model (C). *See Figure 1*.

**Figure 4.**
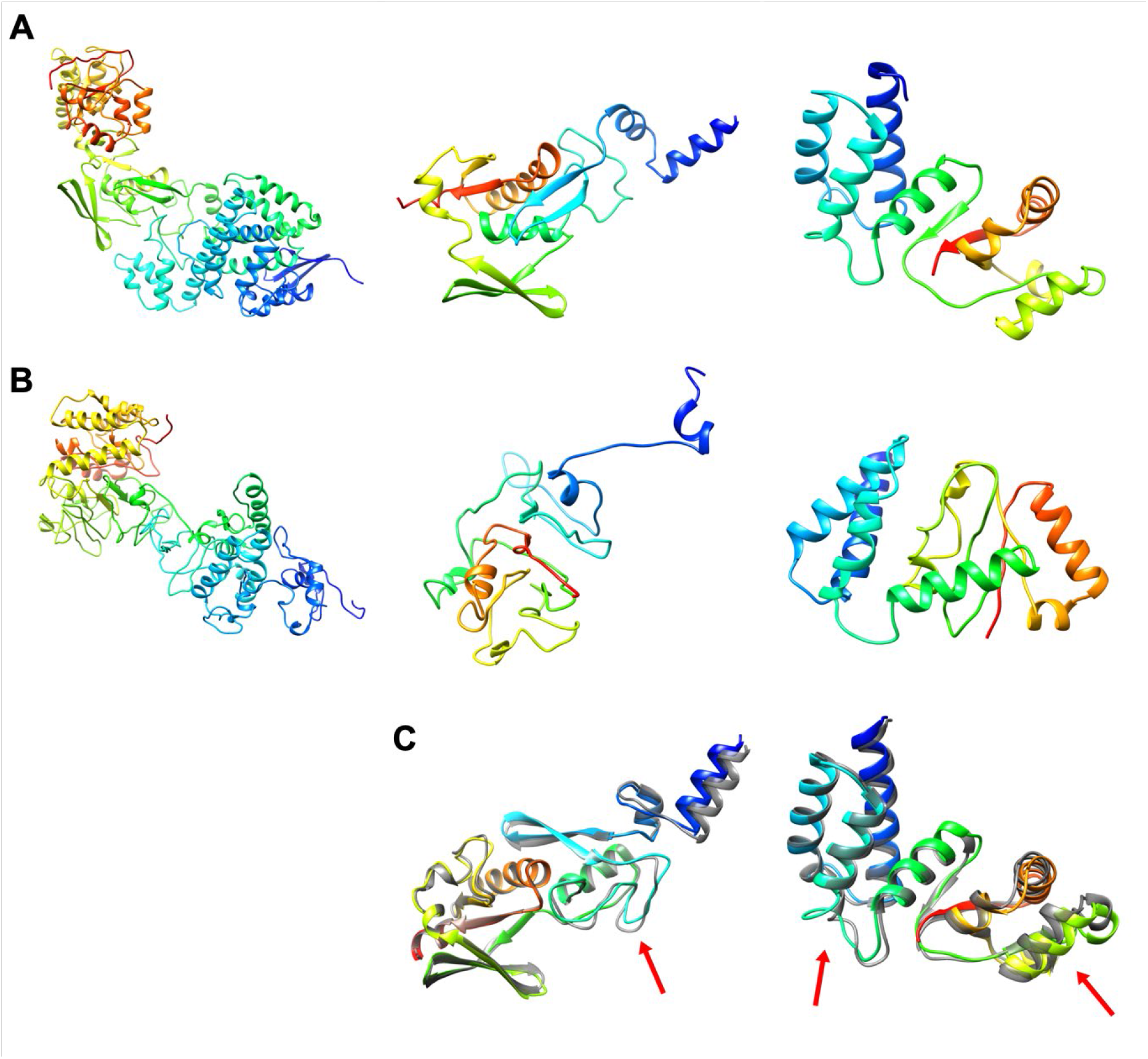
Protein models for Papain-like proteinase (PL-PRO, nsp3): FeigLab (A), Zhang group (B), AlphaFold models and their refined models (C). Domains for residues 1260–1570, 1571–1762, and 1763–1927 are shown left, center, and right, respectively. *See Figure 1*.

**Figure 5.**
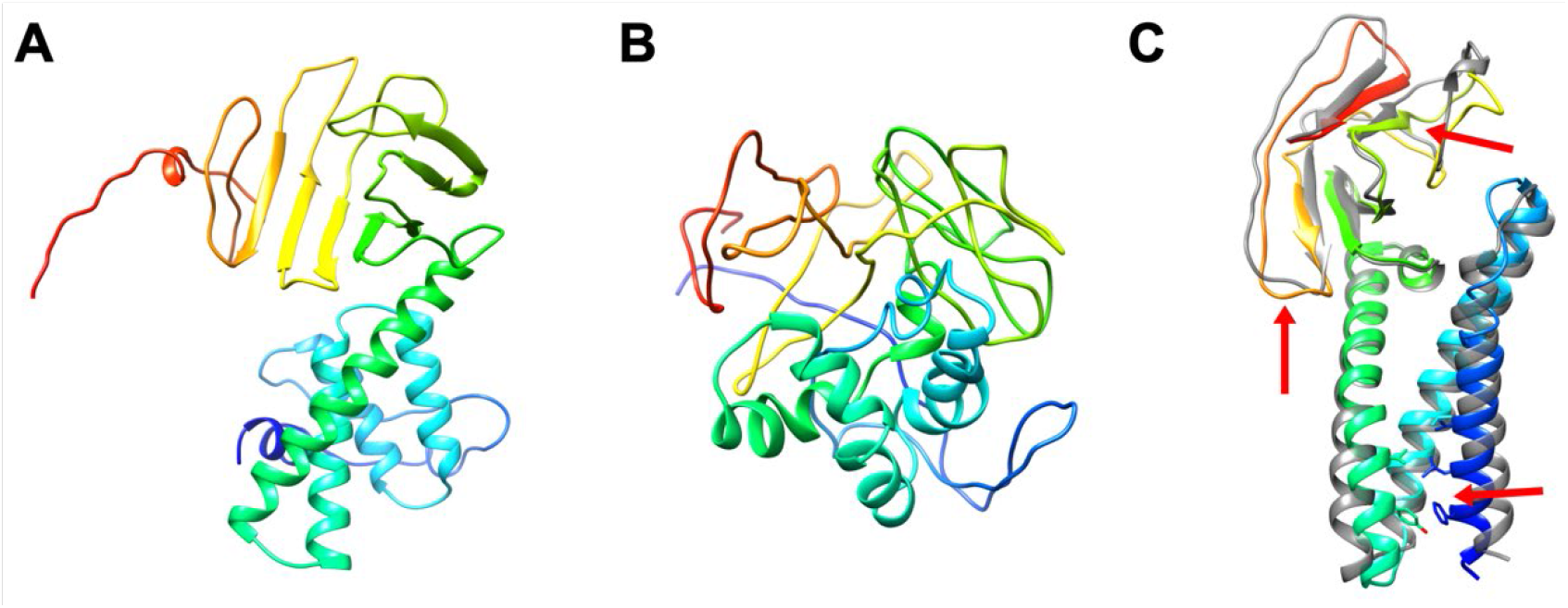
Protein models for ORF3a: FeigLab (A), Zhang group (B), AlphaFold model and its refined model (C). *See Figure 1*.

**Figure 6.**
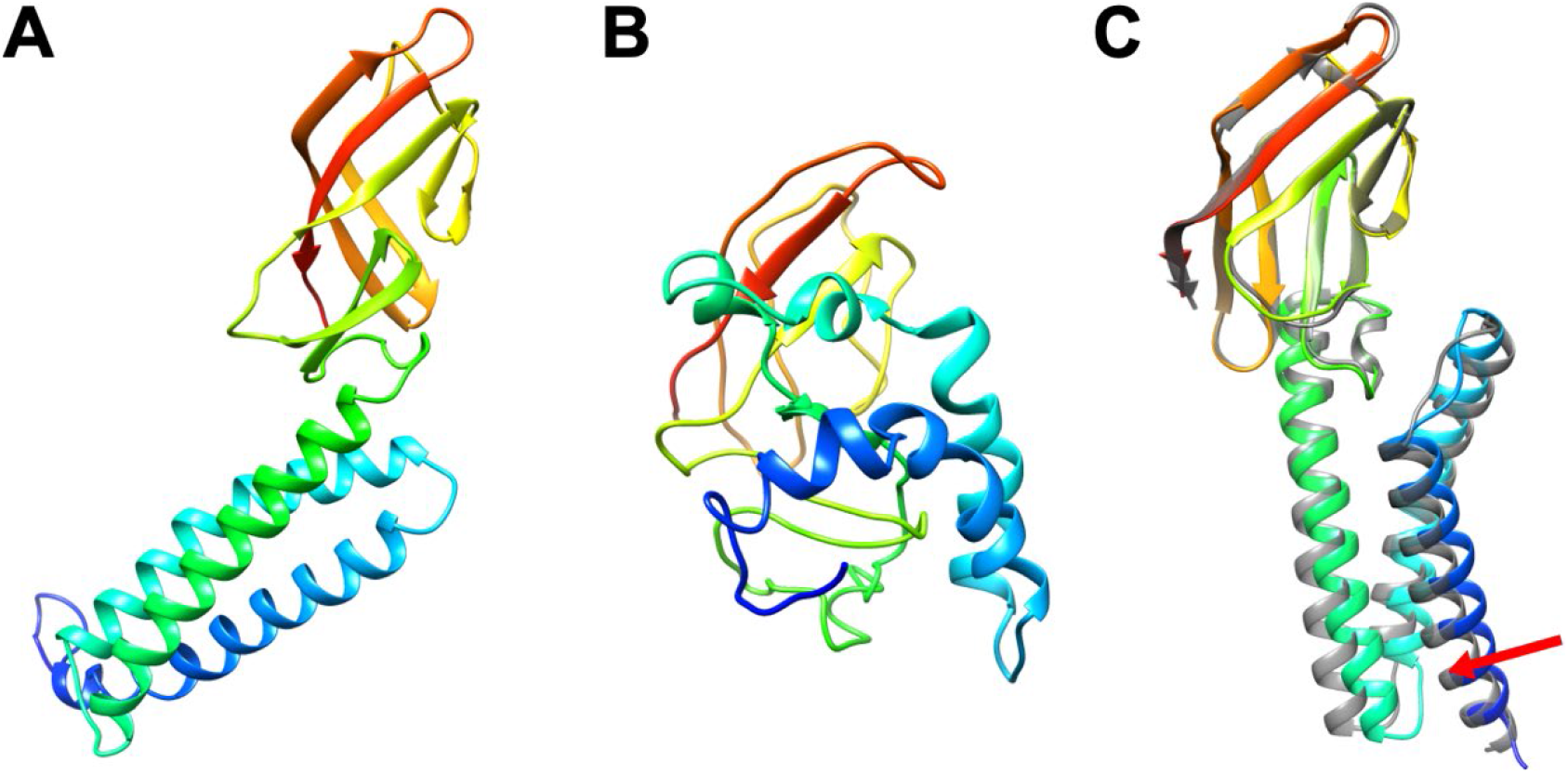
Protein models for Membrane glycoprotein: FeigLab (A), Zhang group (B), AlphaFold model and its refined model (C). *See Figure 1*.

**Figure 7.**
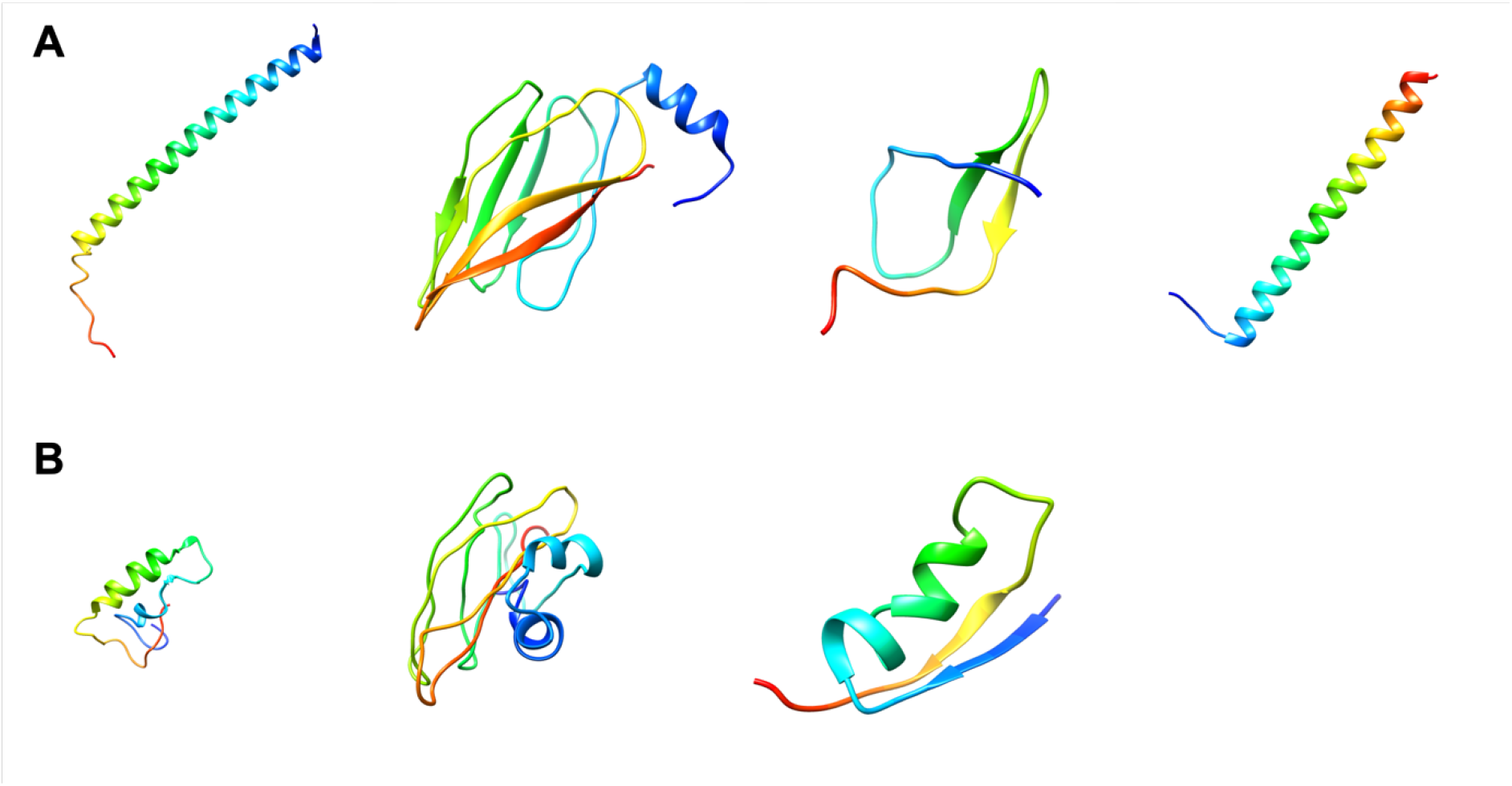
Protein models from FeigLab (A) and Zhang group (B). From the left to the right, protein models for ORF6, ORF8, ORF10, and ORF7b are shown. *See Figure 1.*

We evaluated MolProbity score^11^ for all available models (**Table 3**). Although this score does not indicate whether a model reflects the overall correct structure or not, it can tell if a given model satisfies basic protein stereochemistry or not. All of the models generated from our modeling pipeline has less than 1.5 MolProbity score. Especially, steric clashes and rotamer outliers, rarely exist. Most of the AlphaFold models also have good MolProbity scores, although there are sometimes a few numbers of steric clashes between atoms and some rotamer outliers. After refining those models, most of the poor local geometries could be resolved, and the resulting MolProbity scores after refinement of the AlphaFold models are very good. In contrast to these models, models from Zhang group have poor local geometries as measured by the MolProbity score. These models have numerous atomic clashes, poor side-chain conformations, and bad backbone dihedral angles that generally suggest poor stereochemistry.

**Table 3.** MolProbity scores for the modeled proteins

| Protein name | FeigLab | AlphaFold | Refined AlphaFold | Zhang |
| --- | --- | --- | --- | --- |
| nsp2 | 1.34<br>(0.9 / 0.7% / 89.3%) | 1.68<br>(4.8 / 1.7% / 96.3%) | 0.55<br>(0.1 / 0.4% / 98.2%) | 4.41<br>(281.87 / 7.2% / 68.6%) |
| nsp4 | 1.13<br>(0.3 / 0.9% / 90.6%) | 1.30<br>(3.4 / 1.6% / 98.0%) | 0.73<br>(0.3 / 0.0% / 97.3%) | 4.20<br>(216.46 / 6.6% / 76.1%) |
| nsp6 | 0.94<br>(0.4 / 0.0% / 95.8%) | 1.28<br>(1.8 / 2.9% / 98.2%) | 0.66<br>(0.0 / 0.0% / 97.1%) | 5.12<br>(292.67 / 34.9% / 38.5%) |
| PL-PRO<br>(nsp3) | 1.04<br>(0.5 / 0.5% / 94.6%) | 1.25<br>(2.2 / 2.2% / 98.9%) | 0.77<br>(0.9 / 0.0% / 98.6%) | 4.18<br>(215.81 / 6.5% / 77.2%) |
| ORF3a | 1.20<br>(0.5 / 0.0% / 90.5%) | 2.40<br>(6.0 / 5.1% / 90.7%) | 1.09<br>(0.6 / 0.0% / 94.3%) | 4.10<br>(66.68 / 12.9% / 52.4%) |
| Membrane<br>glycoprotein | 0.74<br>(0.0 / 0.5% / 96.4%) | 1.38<br>(1.9 / 3.7% / 99.0%) | 0.50<br>(0.0 / 0.0% / 98.4%) | 4.55<br>(95.65 / 28.6% / 44.1%) |
| ORF6 | 0.50<br>(0.0 / 0.0% / 98.3%) | N/A | N/A | 4.02<br>(31.79 / 25.0% / 49.2%) |
| ORF8 | 1.38<br>(1.0 / 0.9% / 89.1%) | N/A | N/A | 4.45<br>(86.1 / 24.3% / 45.4%) |
| ORF10 | 1.00<br>(0.0 / 0.0% / 91.7%) | N/A | N/A | 3.50<br>(17.6 / 17.1% / 72.2%) |
| ORF7b | 0.50<br>(0.00 / 0.0% / 100.0%) | N/A | N/A | N/A |

\* Detailed geometric features for MolProbity score, clash score, rotamer outlier, and residues with backbone torsions in Ramachandran-favored regions are shown in parentheses. The clash score is the number of clashes per 1,000 atoms. Rotamer outlier and Ramachandran favored are percentages of residues with rotamer outliers and favored Ramachandran angles, respectively.

## METHODS

Structure models of SARS-CoV-2 proteins were predicted by inter-residue distance prediction-based modeling, followed by molecular dynamics (MD) simulation-based refinement. We used trRosetta^8^ to predict inter-residue distance and orientations and build tertiary structure models. We also used AlphaFold models^4^ as starting models for MD-based refinement.

### Inter-residue distance prediction-based model preparation

We applied the trRosetta method to generate inter-residue distance predictions and to build initial models for further refinement. The original machine-learning trRosetta pipeline was modified to be applied to multiple domain proteins. We iteratively searched sequences and predicted inter-residue distances where contact information was not enough to build a model until all the residues could be built or there was no contact information update. We built 10 models for each protein, and the lowest energy structure was selected for the following refinement step.

In addition to trRosetta-based modeling, we took AlphaFold models from their web page (https://deepmind.com/research/open-source/computational-predictions-of-protein-structures-associated-with-COVID-19) as another set of initial machine-learning based models.

### Molecular dynamics simulation-based refinement

Our latest molecular dynamics simulation-based refinement protocol was applied to the protein models. The method is an improved version of our previous protocol used during CASP13^9^. Generally, we followed our previously published iterative protocol, but without iterations. We ran more (10 trajectories) and longer (200 ns) simulations at 360 K instead of 298 K. At the scoring step, we used RWplus^12^ instead of Rosetta score^13^.

For two of the AlphaFold models, nsp2 and nsp4, the models were split into domains to reduce the computational cost. nsp2 was split into two domains based on its discontinuity: 1–345 and 438–638. nsp4 was split into three domains by visual inspection: 1–273, 274–399, and 400–489.

## ACKNOWLEDGEMENTS

We are grateful to DeepMind team for providing AlphaFold models to the community. This research was supported by National Institutes of Health Grants R35 GM126948. Computational resources were used at the National Science Foundation’s Extreme Science and Engineering Discovery Environment (XSEDE) facilities under Grant TG-MCB090003.

